# Naringenin downregulates inflammation-mediated nitric oxide overproduction and potentiates endogenous antioxidant status during hyperglycemia

**DOI:** 10.1101/2020.04.20.050880

**Authors:** Kanwal Rehman, Ikram Ilahee Khan, Muhammad Sajid Hamid Akash, Komal Jabeen, Kamran Haider, Muhammad Tariq

## Abstract

Nitric oxide (NO) is a key regulating factor for physiological functions, when elevated during inflammatory conditions can lower endogenous antioxidant levels. Increased NO interacts with oxygen or other ROS to generate peroxynitrite, a potent oxidant which induces oxidative stress. Analgesic effects of naringenin (NRN), a flavanone has been demonstrated by inducing anti-inflammatory effects in O_2_^−•^-mediated inflammation. NRN stimulates antioxidant enzymes and also improves glucose uptake. Hence this study was designed to look for therapeutic effects of NRN and in comparison, to metformin (MET) on inflammation-mediated increased NO and decreased antioxidant superoxide dismutase (SOD) in diabetic rat model with compromised glycemic and lipid profile. After single intraperitoneal injection of alloxan (120 mg/kg), the rats were equally divided as Group 1 and 2 which received normal saline and no-treatment respectively while group 3 and 4 received MET 50 mg/kg/day and NRN 50 mg/kg/day respectively. Blood samples were collected at 0, 15^th^ and 30^th^ day of treatment period. Results showed that alloxan significantly increased serum level of glucose (P<0.001), NO (P<0.001) and inflammatory biomarkers (TNF-α, IL-6), however, it expressively decreased serum SOD and insulin level. While, NRN significantly downregulated glucose (P<0.05), lipid profile, TNF-α, IL-6 and normalized level of NO (P<0.01). It also improved SOD level as compared to that of MET-treatment. Histopathology of pancreas also showed significant improvement in morphology after NRN treatment. This work delivers that NRN exerts anti-oxidant effect in part by downregulating the inflammation-mediated NO overproduction and improving level of SOD resulting in potentiation of endogenous antioxidant defense.

## Introduction

Type 2 Diabetes mellitus (DM) is globally spread chronic metabolic disorder that occurs due to the resistance of insulin or high blood glucose level [1]. DM is characterized by high blood sugar level and/or hyperglycemia that causes auto-oxidation of glucose, impairs activity of antioxidant enzymes and may alter glutathione metabolism. All these conditions lead to the increased production of reactive oxygen species (ROS). ROS can cause further damage and dysfunction of pancreatic β-cells. Due to this reason, ROS are among the major causes of DM and its long term complications [2]. Pathophysiology of DM includes insulin resistance owing to decrease uptake of glucose due to which intracellular and extracellular hypoglycemia and hyperglycemia occurs respectively. Both of these conditions can also cause further complications in diabetic patients (Asmat *et al*., 2016).

Nitric oxide (NO) production, key factor of regular endothelial and vascular function is considered to elevate in inflammatory conditions with lower endogenous antioxidant levels. This increased NO in association with other generated ROS contributes for the induction of oxidative stress. It has been reported that during any inflammatory condition, the release of inflammatory cytokines, such as IL-6 and TNF-α can cause overproduction of NO via potentiating inducible nitric oxide synthetase activity at a higher rate [3]. Nitric oxide contains unpaired electron which plays a considerable role in different signaling mechanisms [4]. NO interacts with oxygen or other free radicals and generates a potent oxidant, peroxynitrite causing various other disease complications [5]. Superoxide anion O_2_^−•^ produce peroxynitrite by interacting with NO however, superoxide dismutase (SOD), an antioxidant enzyme, coverts O_2_^−•^ into hydrogen peroxide [6]. Hence, numerous approaches can be utilized to avert or restore peroxynitrite-induced cellular damage. One strategy can be reduction in NO production or other can be increasing the level of SOD as it declines the formation of free radical O_2_^−•^-mediated peroxynitrite thus reduces oxidative stress. It has also been reported that hyperglycemia might be sometimes associated with an increased NO level [7] which may contribute to the development of DM-related vascular diseases. As far as the treatment of DM is concerned, [7], metformin is a widely used antidiabetic drug which works in different ways including improved muscle glucose uptake and reduction in the absorption of glucose from the small intestine into circulatory system [8].

Recently, flavonoids, the group of compounds with a vast range of biological activities often present in herbs, vegetables and fruits are gaining attention for their therapeutic efficacy against complicated chronic diseases. There are different types of flavonoids which includes flavanones, isoflavones, flavanols. Naringenin (NRN) belongs to the flavanone class of flavonoids and is usually present in grape fruits or citrus fruits etc. [9]. It has been reported that NRN is abundantly present mostly in citrus fruits, like in grapefruit (87 mg/200 mL), in orange juice (4.26 mg/200 mL), and to a lesser extent in lemon juice (0.76 mg/200 mL). NRN has been observed safe at high doses even at 5000 mg/kg and has good anti-hyperglycemic properties. The proposed mechanism behind NRN therapeutic activity particularly against reactive species is due to the presence of hydroxyl groups in its ring which may indulge the free radicals generated due to hyperglycemia [10]. Interestingly, glucose uptake in primary porcine myotubes has been improved after exposure to NRN [11]. Moreover, the improved glucose uptake has also been associated with insulin-mediated sensitivity in L6 myotubes rendered insulin resistant via exposure to palmitate. Further, GLUT4 translocation was also observed in these myotubes after treatment with NRN (50 μM and 75 μM) for 16 h [12]. These studies clearly support the antidiabetic potential of NRN. In addition to this, NRN has also been reported for its anti-inflammatory potential against sodium-induced colitis model [13]. Analgesic effects of NRN has also been observed to demonstrate by inducing anti-inflammatory effects in O_2_^−•^-mediated inflammatory pain model [14]. Moreover, NRN has shown to potentiate the antioxidant status by stimulating activities of antioxidant enzymes including SOD to mitigate streptozotocin-induced liver injury [15].

Hence, taking in to account the antihyperglycemic, anti-inflammatory and antioxidant potential of NRN, we tried to emphasize on comparable healing strategy, involving the utilization of natural product being more efficient with minimum toxicity and to mark the foremost pathways responsible for regulating the typical glucose absorption. We investigated the anti-inflammatory and anti-oxidant roles of NRN against alloxan-induced diabetic animal model with compromised antioxidant status and inflammation-mediated NO elevated level. To the best of our knowledge, till date, the effect of NRN alone and in comparison, to the effects of MET on serum NO and antioxidant enzyme SOD during hyperglycemic state has not been elucidated.

## Material and methods

### Chemicals

Alloxan (Alfa Aesar™ by Thermo Fisher Scientific), metformin (Acros Organics by Thermo Fisher Scientific), Naringenin (Alfa Aesar™ by Thermo Fisher Scientific) and all other chemicals used were of analytical grade.

### Study design

About Twenty-four white albino Wistar rats with weight between 180-250 g were purchased and kept in the animal house of University of Agriculture Faisalabad (UAF), Pakistan and housed at ambient temperature (25 ± 5 °C). ambient temperature (25 ± 5 °C). All experimental procedures were carried out at UAF, in accordance with the approved laboratory animal Bio-safety guidelines, Biosafety/Bioethics protocol of Institutional Biosafety committee (IBC) and Bioethics committee and research board DGS/3601-04). The guidelines were used for appropriate animal housing for which all rats were fed on normal diet with water *ad libitum* and were allowed to acclimatize for two weeks before the start of experiments, proper blood sampling was done that is about 2.0-2.5 ml blood samples were attained by tail vein method from each rat of every group and animal pain was minimized by adopting merciful killing of rats at the end of study period as rats were sacrificed by cervical dislocation and the abdomen was dissected, for the removal of required tissues for histopathological analysis. According to study protocol, diabetes was induced in all rats by injecting a single intraperitoneal injection of alloxan (120 mg/kg). After 3-4 days, induction of diabetes was confirmed by monitoring the fasting/random blood glucose level. Later on, the rats were equally divided into four groups each containing 6 rats in individual cages for 30 days according to the study protocols. Group 1 was marked as normal control group (**NC**). Group 2 was marked as diabetic control (**DC**) group in which no therapeutic intervention was made throughout the study period. Further, Group 3 was marked as standard treated group, this group was treated with metformin (**MET**) as standard drug with a dose of 50 mg/kg/day. Likewise, group 4 was marked as naringenin treated group (**NRN**) as exposed to therapeutic intervention of NRN (50 mg/kg/day).

### Blood sampling

Blood samples were collected at 0 day i.e. before starting the treatment. At 15^th^ and 30^th^ day of the study period, blood samples were again collected for measuring the required biochemical parameters. The collected blood samples were centrifuged at 3000 RPM for 20 minutes and the serum was stored at −20 °C till further biochemical analysis.

### Biochemical analysis

#### Estimation of glycemic biomarkers

To estimate the effect of treatment on hyperglycemia, we measured the blood glucose and insulin level before, during and at the end of treatment period using their respective assay reagent kits.

#### Estimation of serum liver and kidney biomarkers

To evaluate any toxic effect of MET and NRN treatment on liver and/or kidney of alloxan-induced diabetic rat model, liver function biomarkers including alanine aminotransferase (ALT), alkaline phosphatase (ALP) and aspartate aminotransferase (AST) and kidney function biomarkers notably, urea and creatinine were estimated before, during and at the end of treatment using respective assay reagent kits.

#### Estimation of lipid biomarkers

Likewise, to investigate the effect of therapeutic treatment on lipid profile of alloxan-induced diabetic rat model, we estimated the serum level of lipid biomarkers namely high-density lipoprotein (HDL), low density lipoprotein (LDL) and triglycerides (TGs) before, during and the end of treatment using respective assay reagent kits.

#### Estimation of serum anti-oxidant and nitric oxide

Similarly, we measured the serum level of superoxide dismutase (SOD), an antioxidant enzyme to investigate the antioxidant properties of therapeutic intervention used in this study against alloxan-induced diabetic rat model. We measured the level of anti-oxidant by calorimetric method with commercially available kits at the start, mid and end of treatment.

#### Estimation of serum nitric oxide

We used commercially available kit to investigate the effect of NRN treatment on serum NO level of alloxan-induced diabetic rat model at beginning, mid and end of study,

### Histopathological analysis

At the end of treatment period, At the end of treatment, rats were sacrificed by cervical dislocation and the abdomen was dissected, pancreas, were removed for histopathological analysis. Tissue sections from the pancreas were prepared by fixation and sectioning followed by staining for histopathological analysis.

## Results

### Effect of treatment on glycemia

The efficacy of NRN against elevated level of blood glucose and decreased insulin secretion in alloxan-induced diabetic rats was assessed. The effect of treatment is shown in figure 1, (A) glucose and (B) insulin. After alloxan exposure, we found significant increase (P<0.001) in the serum level of glucose and reduced insulin secretion when compared with that of non-diabetic control group. However, at the 15^th^ day of treatment period, MET treated group showed decreased (P<0.001) glucose level and improved insulin secretion as compared to that of DC group. However, this pattern of improved glycemic profile was more significant (P<0.05) after exposure to NRN when compared with DC and MET-treated group at the end of treatment period. Similarly, in case of effect of treatment on serum insulin, we found that due to the IP injection of alloxan, the serum level of insulin was significantly decreased as compared to that of normal rats (Figure 1B) but once, the treatment was started, we found that NRN significantly increased the serum level of insulin even in a better way when it was compared with that of MET-treated rats at the end of treatment period (Figure 1B).

**Figure 1:**
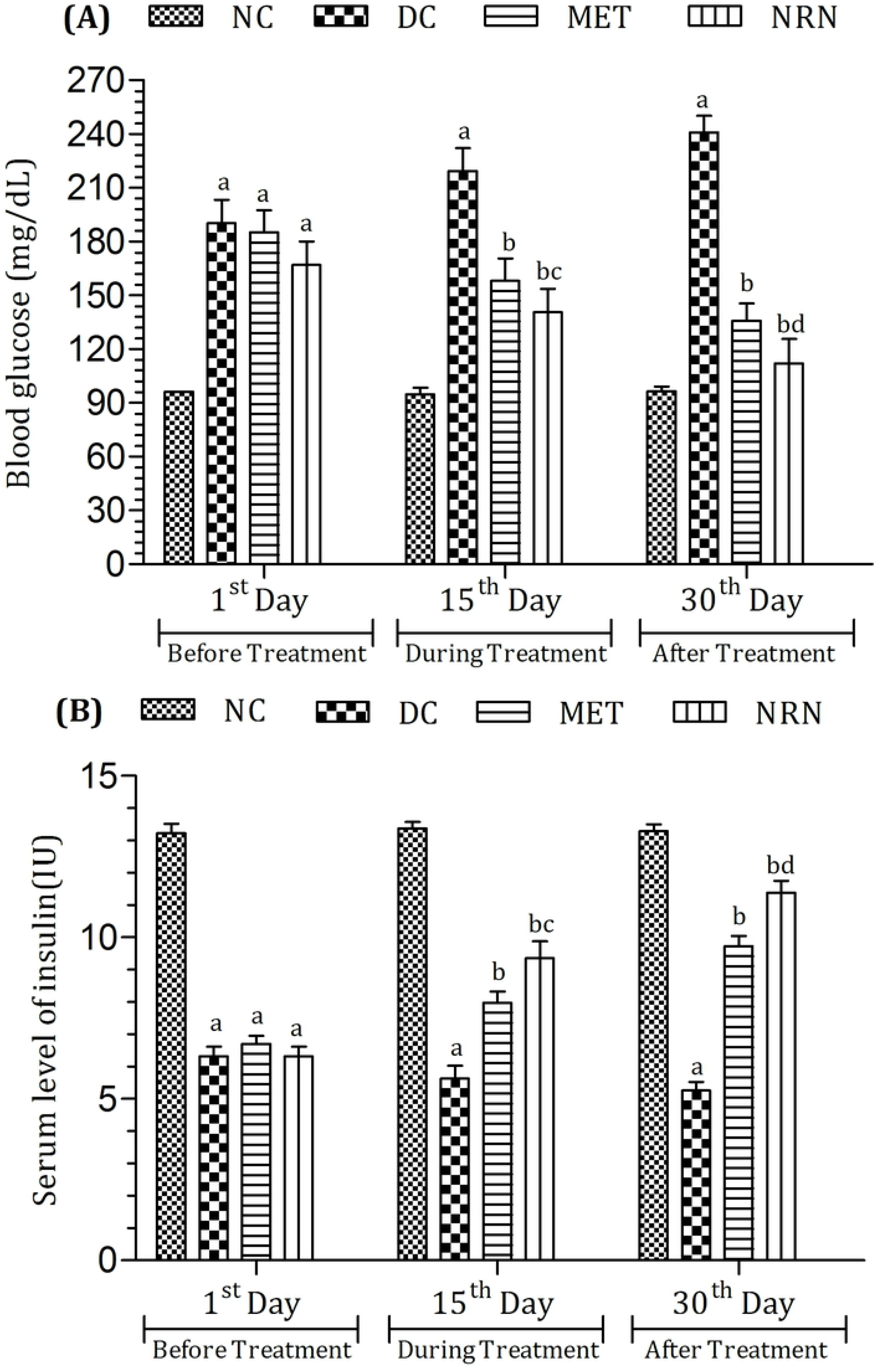
Effect of treatment on (A) glucose and (B) insulin. To estimate the effect of treatment on glycemia, we measured the serum level of glucose and insulin at 1^st^, 15^th^ and 30^th^ day of the treatment period. The level of significant difference was estimated by Bonferroni post-test using two-way ANOVA. ^a^ represent P<0.001 when compared with CN-group. ^b^ represents P<0.001 when compared with DC-group. ^c^ represents P<0.05 when compared with MET-group. ^d^ represents P<0.001 when compared with MET-group.

### Effect of treatment on lipidemia

Similarly, NRN was also evaluated for its effect on the lipid profile including serum level of HDL, LDL and TGs in alloxan-induced diabetic rats before and after treatment in comparison to non-treated DC and MET-treated groups. The effects are shown in figure 2, (A) HDL, (B) cholesterol, (C) TGs and (D) LDL. Before the start of therapeutic intervention, it was observed that alloxan significantly increased (P<0.001) the serum level of all above parameters except HDL in rats as compared with that of non-treated normal control group. However, at 15^th^ day of treatment, both MET- and NRN-treated groups significantly (P<0.001) showed improvement in the serum level of HDL (Figure 2A). Further, at the end of treatment, NRN-treated group had a significant improvement (P<0.001) in the serum lipid profile with an evident increase in the level of HDL as compared with MET and DC-treated group (Figure 2).

**Figure 2:**
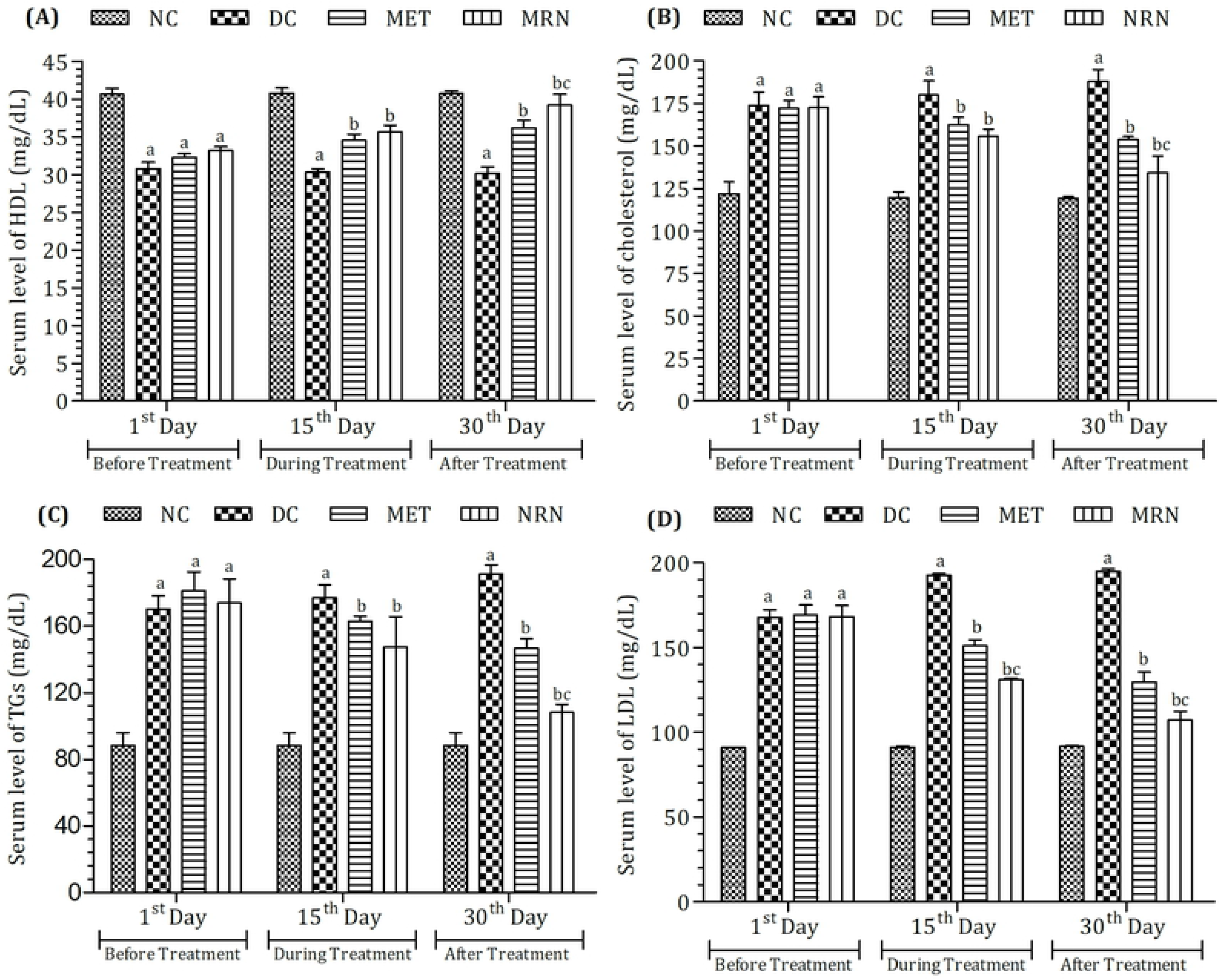
Effect of treatment on (A) HDL, (B) cholesterol, (C) TGs and (D) LDL. To estimate the effect of treatment on lipid profile, we measured the serum level of HDL, cholesterol, LDL and TGs at 1^st^, 15^th^ and 30^th^ day of the treatment period. The level of significant difference was estimated by Bonferroni post-test using two-way ANOVA. ^a^ represent P<0.001 when compared with CN-group. ^b^ represents P<0.001 when compared with DC-group. ^c^ represents P<0.001 when compared with MET-group.

### Effect of treatment on liver function enzymes

The liver function enzymes including serum level of AST, ALT and ALP were assessed to evaluate the effect of treatment on alloxan-induced diabetic rats. As depicted in figure 3, (A) AST, (B) ALT and (C) ALP, the serum level of all enzymes after alloxan induction started to increase (P<0.001) as compared to that in NC group (Figure 3). However, after starting the therapeutic intervention, NRN and MET showed the improvement in these enzymes level. A significant (P<0.001) decline in the levels of AST, ALT and ALP was observed at the end of study with both MET and NRN groups, where NRN showed more efficacy as compared to that of MET group.

**Figure 3:**
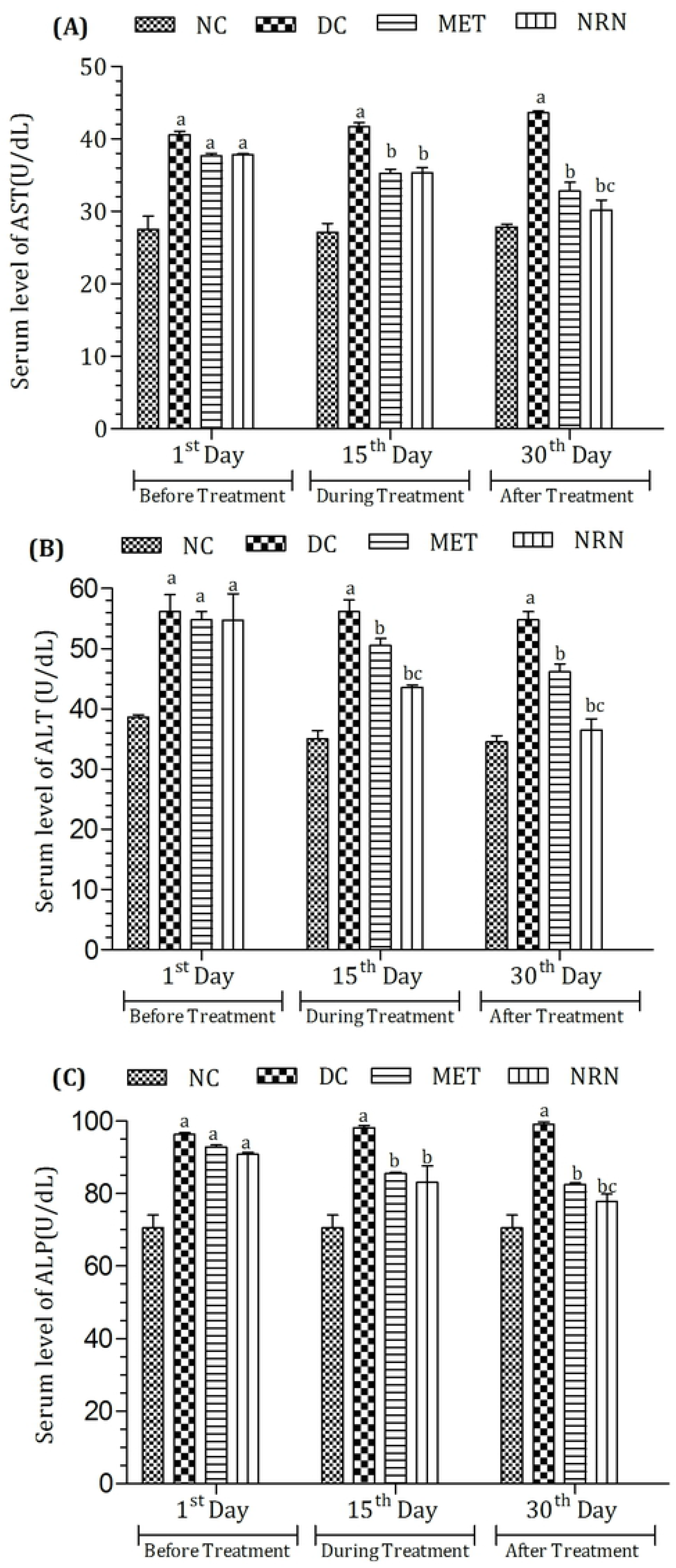
Effect of treatment on (A) AST, (B) ALT and (C) ALP. To estimate the effect of treatment on liver function biomarkers, we measured the serum level of AST, ALT and ALP at 1^st^, 15^th^ and 30^th^ day of the treatment. The level of significant difference was estimated by Bonferroni post-test using two-way ANOVA. ^a^ represent P<0.001 when compared with CN-group. ^b^ represents P<0.001 when compared with DC-group. ^c^ represents P<0.001 when compared with MET-group.

### Effect of treatment on kidney biomarkers

As shown in figure 4, the serum urea and creatinine as kidney biomarkers followed the same pattern of increase as liver function enzymes particularly after the induction of alloxan. However, NRN-treatment significantly (P<0.01) declined the serum level of kidney biomarkers when compared with MET and DC group (Figure 4).

**Figure 4:**
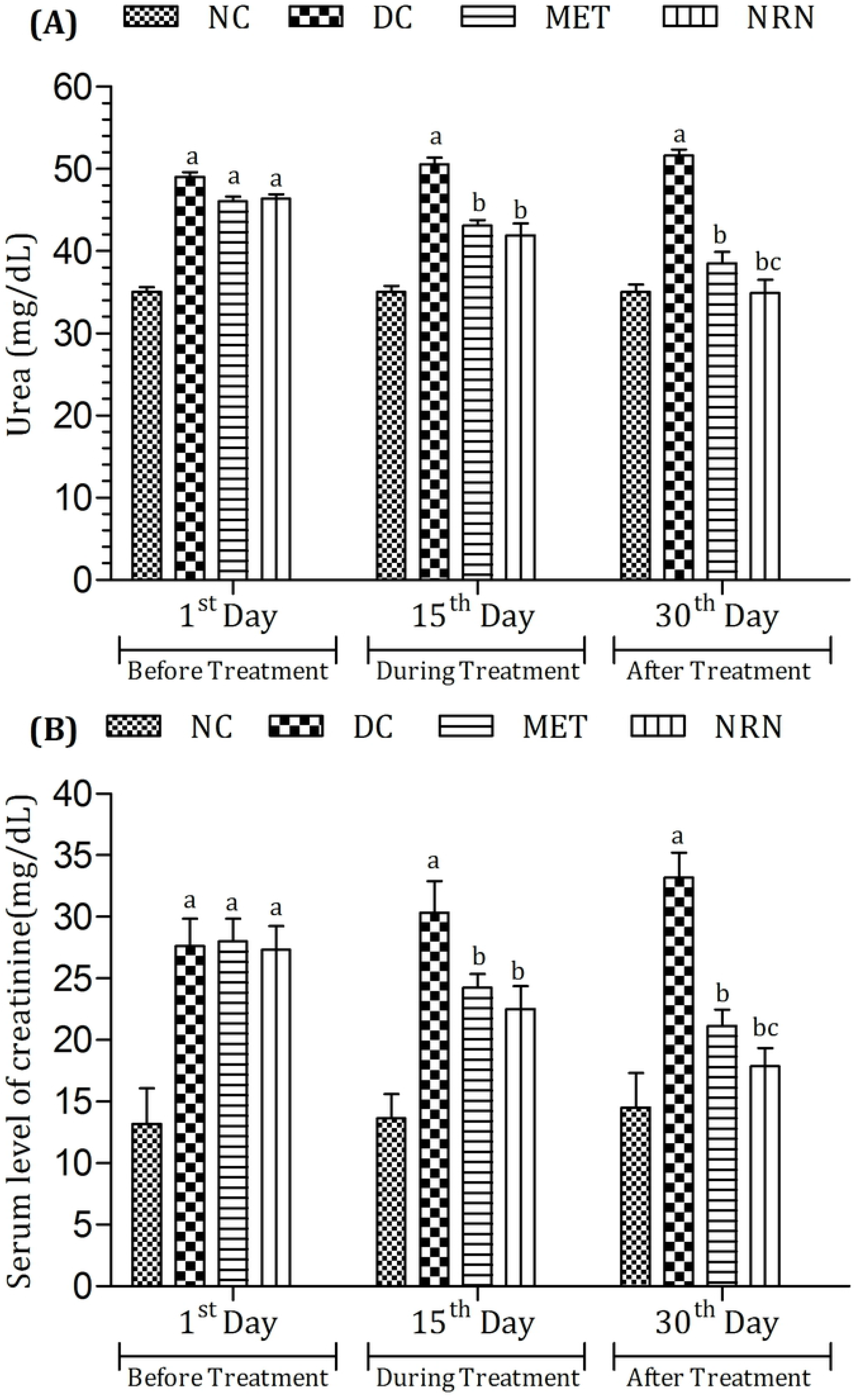
Effect of treatment on (A) Urea and (B) creatinine. To estimate the effect of treatment on kidney function biomarkers, we measured the serum level of urea and creatinine at 1^st^, 15^th^ and 30^th^ day of the treatment period. The level of significant difference was estimated by Bonferroni post-test using two-way ANOVA. ^a^ represent P<0.001 when compared with CN-group. ^b^ represents P<0.001 when compared with DC-group. ^c^ represents P<0.01 when compared with MET-group.

### Effect of treatment on antioxidant status

The antioxidant status including the level of serum SOD and NO was also measured in order to investigate the effect of NRN-treatment on NO and oxidative stress induced by alloxan toxicity as shown in figure 5. At 15^th^ day of treatment, both MET and NRN treated groups significantly (P<0.01) decreased the serum level of NO and improved SOD level when compared as compared to that of DC group with a more pronounced effect of NRN on the same profile at the end of study period showing significant increase in SOD level when compared with MET and DC group indicating the antioxidant property of NRN (Figure 5).

**Figure 5:**
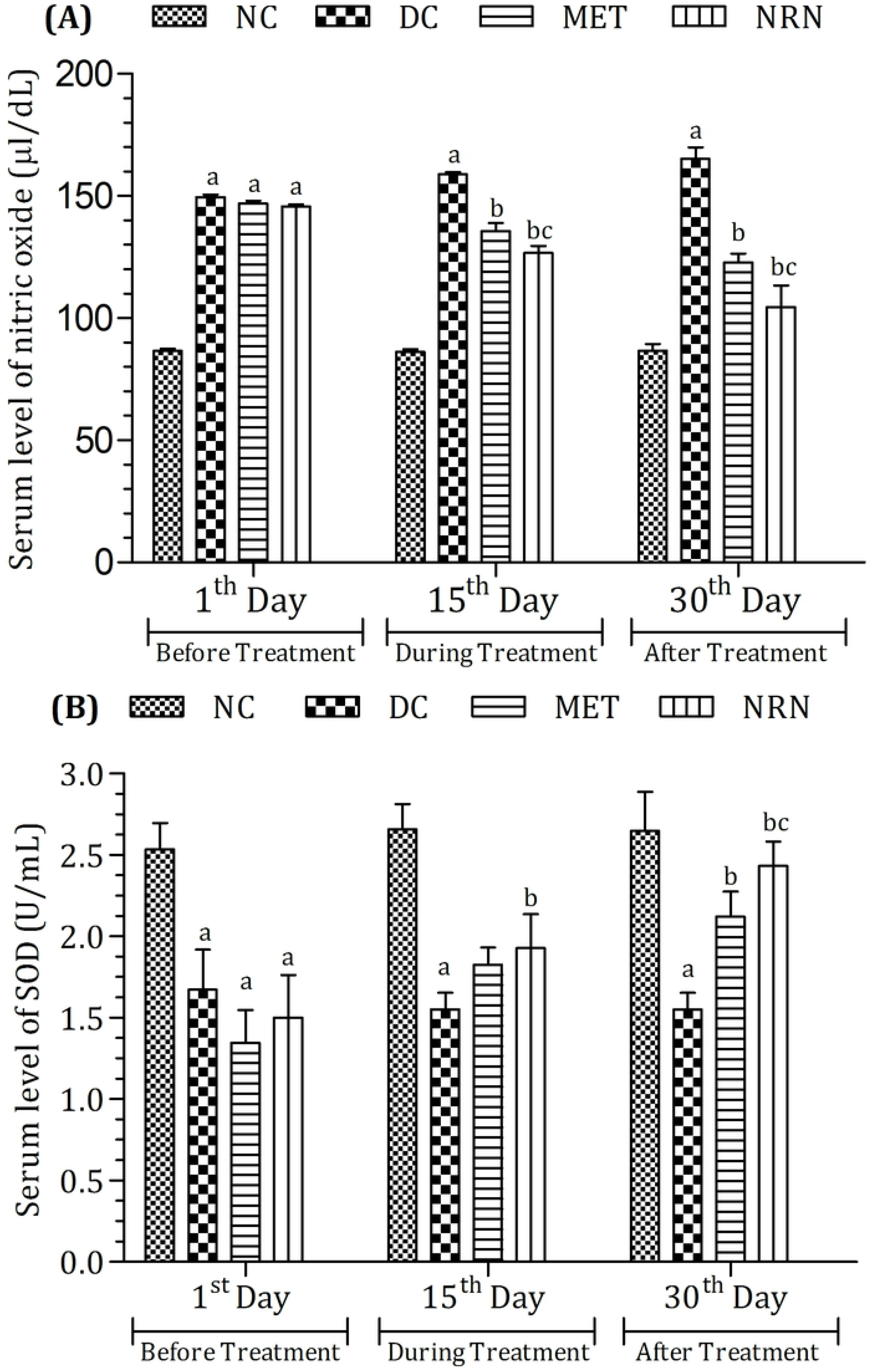
Effect of treatment on (A) nitric oxide and (B) SOD. To estimate the effect of treatment on oxidative stress, we measured the serum level of nitric oxide and SOD at 1^st^, 15^th^ and 30^th^ day of the treatment period. The level of significant difference was estimated by Bonferroni post-test using two-way ANOVA. ^a^ represent P<0.001 when compared with NC-group. ^b^ represents P<0.01 when compared with DC-group. ^c^ represents P<0.05 when compared with MET-group.

### Effect of treatment on proinflammatory mediators

Proinflammatory mediators including serum IL-6 and TNF-α was also measured in order to investigate the role of NRN treatment on regulating the inflammatory responses induced by alloxan (Figure 6). At 15^th^ day of treatment both MET and NRN treated groups significantly (P<0.01) decreased the serum level of IL-6 and TNF-α when compared with DC group. A more pronounced effect of NRN on the same profile at the end of study period showed a significant decrease in IL-6 and TNF-α level when compared with MET group indicating the anti-inflammatory property of NRN (Figure 6).

**Figure 6:**
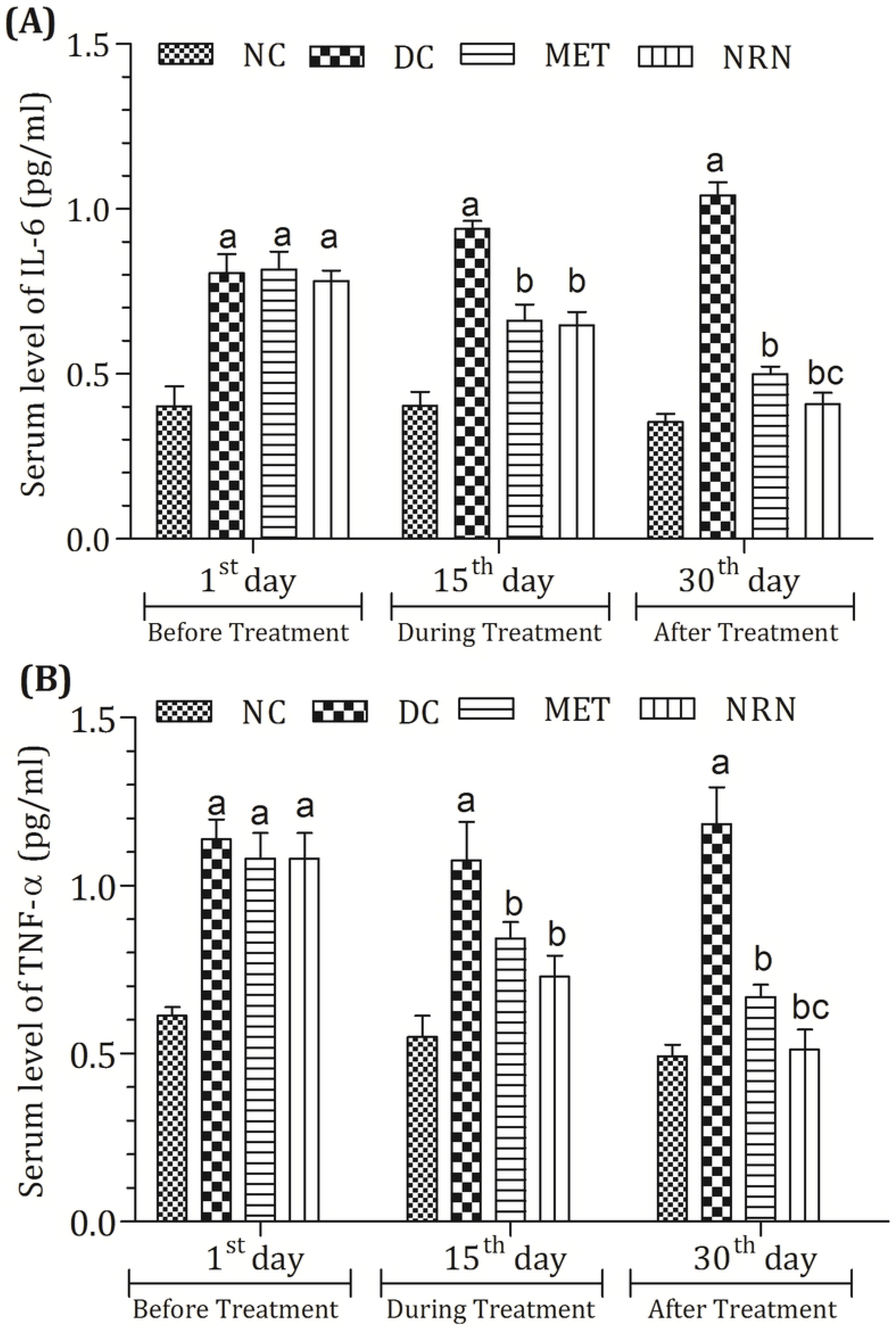
Effect of treatment on (A) IL-6 and (B) TNF-α. To estimate the effect of treatment on proinflammatory cytokines, we measured the serum level of IL-6 and TNF-α at 1^st^, 15^th^ and 30^th^ day of the treatment period. The level of significant difference was estimated by Bonferroni posttest using two-way ANOVA. ^a^ represent P<0.001 when compared with NC-group. ^b^ represents P<0.01 when compared with DC-group. ^c^ represents P<0.05 when compared with MET-group.

### Effect of treatment of histopathology of pancreas

Normal histological architecture of islets of Langerhans was seen in NC group. Moreover, the acinar cells were well arranged. However, the alloxan to rats critically damaged the β-cells of pancreas. In addition, the treatment with MET, or NRN alone showed little damage to the β-cell through or deprived of fractional reinstatement (Figure 7).

**Figure 7:**
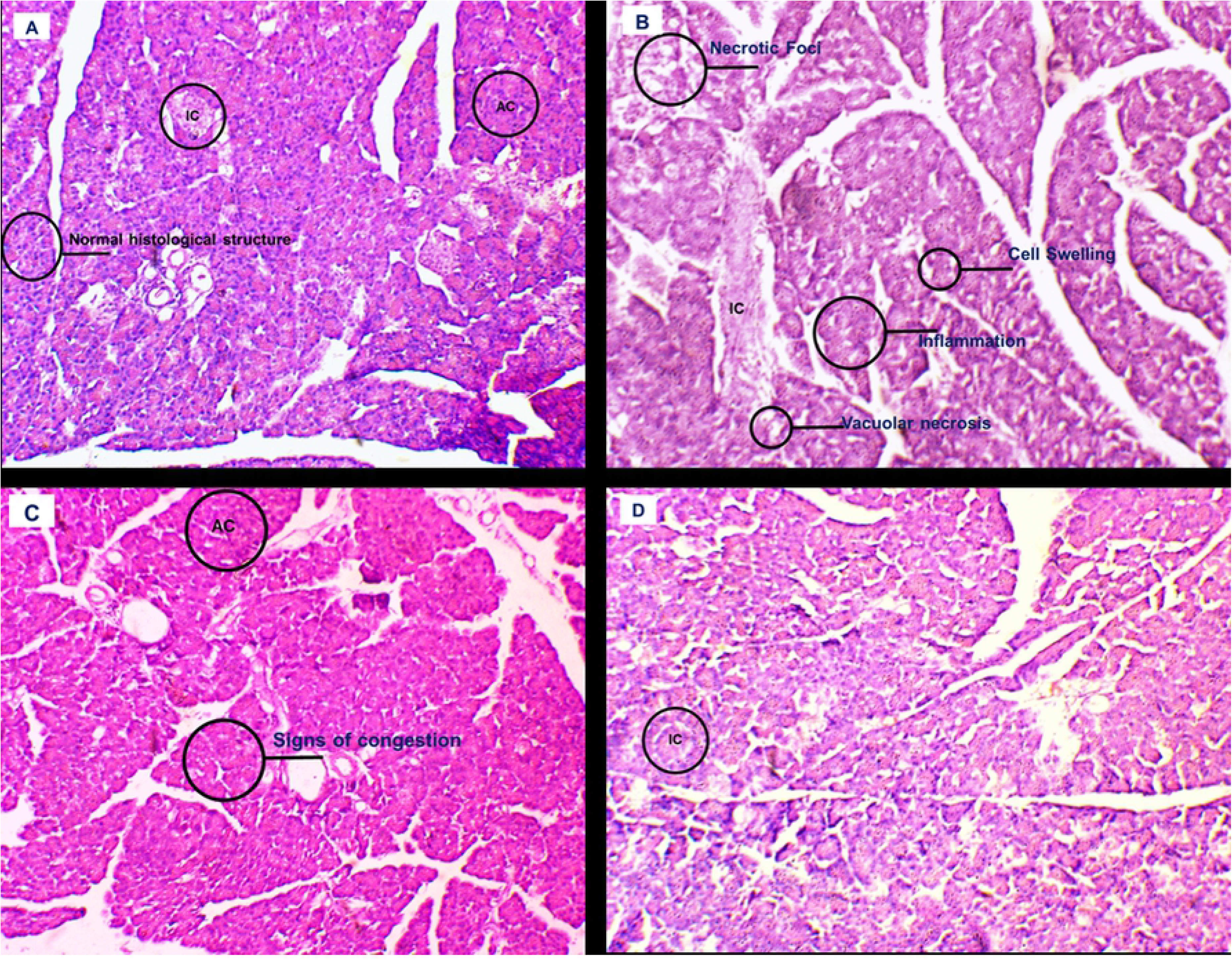
Effect of treatment of histopathology of pancreas: NC-group (A); β-cells appeared normal in numbers. The acinar cells were stained and appeared normal in structure. DC-group (B); β-cells appearance was not normal and there was increased in necrotic foci, cell swelling, inflammation and vacuolar necrosis evident by photomicrography with blood vessels at few places. MET-group (C); the acinar cells were stained and appeared normal in structure. Congestion was present at few places. NRN-group (D); acinar cells were normal in appearance with normally distributed β-cells and blood vessels.

## Discussion

Diabetes mellitus is very common disease of the endocrine system globally, a huge percentage of population is suffering from diabetes every year. According to international diabetes federation, diabetes may affect 300 million of global population till 2025. Likewise, DM has been reported to be among major causes of deaths in developing countries like Pakistan. The effect of NO is considered to have multiple aspects in terms of molecular functioning intracellularly for proper cellular function as well as playing harmful effects during progression of many chronic diseases probably by generation of ROS [16]. Augmented level of serum NO in T2DM has been observed by many researchers in previous works with debatable results and outcomes [17,18]. Increased level of serum NO in patients with DM not only seen responsible for causing DM but has also considered to play a role in DM-associated complications [19]. Irregular NO metabolism may have a role in the pathogenesis of DM. Inhibition of inducible NOS may be of worth for the avoidance of DM-associated microvascular complications. The level of blood glucose and NO is considered sometimes to be directly proportional to each other. Moreover, hyperglycemia can activate the endothelial cells for the release of nitric oxide and its raised level in blood that may cause increased production of RNS [7]. NO is known to raise during inflammatory conditions including metabolic disorders. In addition, it also effects the working of antioxidant enzymes as it interacts with free radical species to generate potent oxidants promoting oxidative stress. When inflammation occurs, the pro-inflammatory mediators like IL-6 and TNF-α affect the working of NOS activity and cause over production of NO [3]. Unpaired electron is possessed by NO molecule [4], therefore, NO interacts with oxygen, produces peroxynitrite and may lead to various other diseases and disorders [5], however, superoxide dismutase (SOD), an antioxidant enzymes, coverts O_2_^−•^ into hydrogen peroxide [6]. Henceforth, many different ways can be adopted to avoid or reinstate peroxynitrite-induced stress and damages. One approach is to slow down the production of NO or to potentiate the antioxidant enzymes so it may help converting free reactive oxygen radical O_2_^−•^ into less reactive H_2_O_2_. A schematic representation of mechanism of action of nitric oxide and NRN has been represented in figure 8.

**Figure 8:**
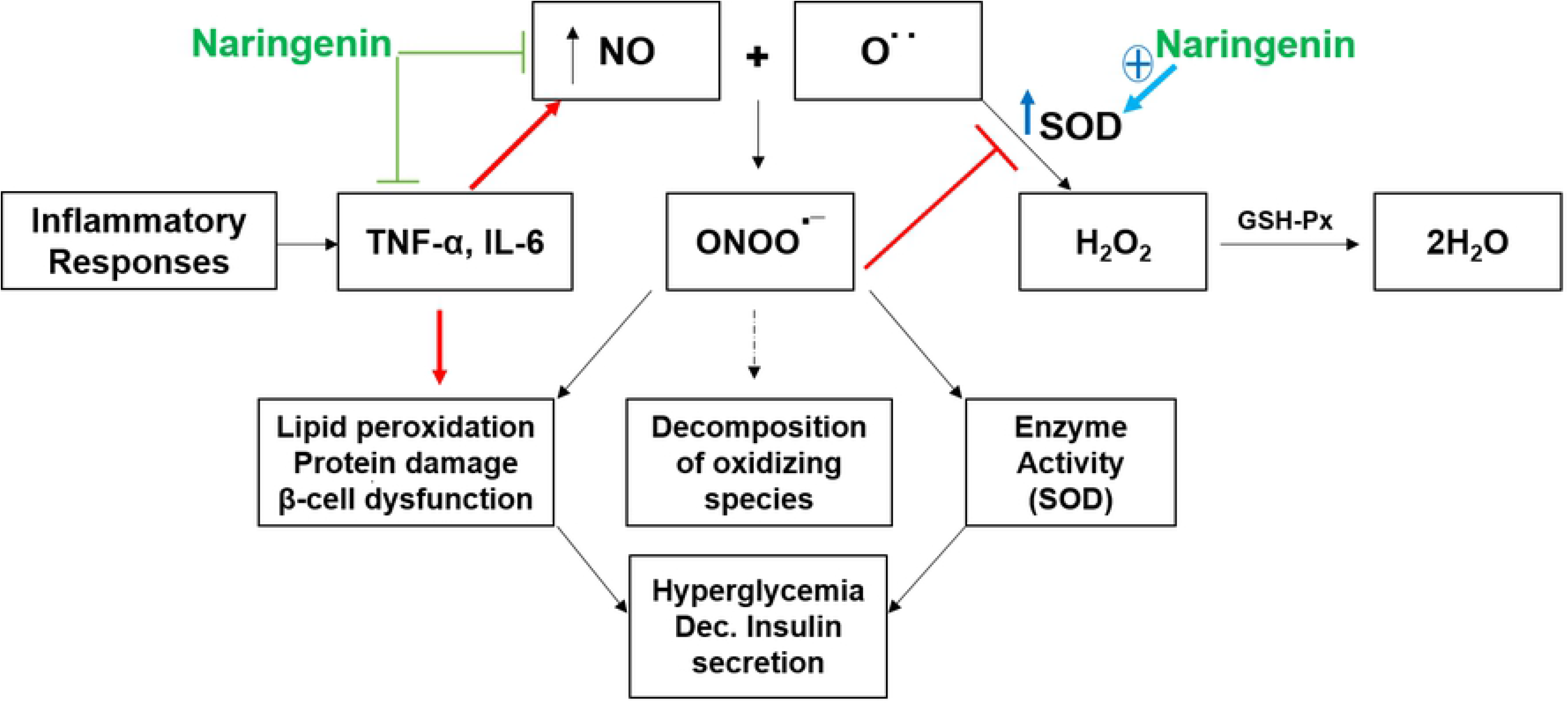
A schematic representation of role of nitric oxide, oxidants, antioxidant SOD, and mechanism of action NRN. During inflammation, the release of inflammatory cytokines, such as interleukins and TNF-α can cause over production of NO via potentiating inducible nitric oxide synthetase activity. This excess NO interacts with superoxide anion O2−• being a free oxygen radical and produces peroxynitrite that may lead to various conditions like lipid peroxidation, oxidative stress, protein damage hyperglycemia and insulin resistance however, superoxide dismutase (SOD), an antioxidant enzymes, coverts O2−• into hydrogen peroxide (H2O2) which is further neutralized by the action of other antioxidant enzymes.

In the present study, we were concerned to elaborate any probable alteration in the serum levels of NO during induction and progression of hyperglycemia in alloxan-induced diabetic rat model after induction of inflammation as evident by elevated levels of IL-6 and TNF-α. These proinflammatory cytokines along with generated reactive NO may cause suppression of antioxidant defense level and induce oxidative stress. For this we also evaluated the level of SOD and compared all these effects with altered biochemical parameter including glycemic, lipid profiles and liver kidney function biomarkers.

NRN is a flavonoid that belongs to the flavanone class of the flavonoids. It is usually present in grapes and citrus fruits etc. NRN has been considered safe at high dose possessing anti-hyperglycemic effect [10]. In hyperglycemia, the glucose intolerance and insulin resistance occurs, both these factors causes increase in blood glucose level. In our study, we investigated the effect of NRN on blood glucose level and serum insulin level as shown in Fig. 1A and B. These figures clearly explained the hypoglycemic property of NRN trough rapid decline in the level of glucose when compared to MET and DC groups. These results support the findings of other studies which exhibit the antihyperglycemic potential of NRN [20,21]. The interesting part in our study is that a much lower dose of NRN (50 mg/kg/day) has produced similar antiglycemic effects in alloxan-induced diabetic rats as compared to the one observed with a greater dose of NRN (75 mg/kg) against alloxan induced diabetic mice [22].

Diabetes and other metabolic disorders are also known to involve in dyslipidemia [23]. So we evaluated the lipid profile before and after treatment in diabetic rats. Interestingly, when we started the treatment with NRN, the HDL level was significantly improved (Fig 2A) while there was significant decrease in the level of cholesterol, triglycerides and LDL (Fig. 2B, C and D) as compared with MET treatment. Our results are comparable to those reported by Assini and his colleagues where mice animal models exposed to NRN treatment (50 mg/kg/day) showed great improvement in lipid profile and tempered hepatic steatosis because of increased cholesterol ingestion [24]. NRN, has also been reported for its ability to reduce synthesis of fatty acid [25] that may probably be a route for regulating lipid profile in diabetic rats [26].

The effect of NRN on alloxan-induced variations in the levels of AST, ALT, ALP (Fig. 3A, B and C) in liver of control and experimental rats was also observed in the current study. NRN, as compared to MET treatment, provided a better prevention against hepatotoxicity in alloxan exposed rats. NRN has been evaluated against various causative factors increasing the levels of these enzymes in different in vivo and in vitro studies [27]. Similar, we investigated levels of kidney function markers namely; blood urea nitrogen (BUN) and creatinine (Fig. 4) which were found to be disturbed after exposure to alloxan. However, significant improvement has been observed after treatment particularly with NRN. This was found to be in accordance with studies reporting renal protective role of NRN probably because of its therapeutic potential to regulate renin angiotensin system [28] or anti-inflammatory efficacy [29,30]. Moreover, a similar dose of NRN (50 mg/kg) for only 2 days exposure has also been claimed to show reduction in lipid peroxidation levels of liver and kidney tissues in alloxan-induced diabetic mice (Sirovina et al 2016).

Moreover, the antioxidant potential of NRN can be considered accountable for the hepato- and renal protective potential of NRN [31] along with regulation of lipid and glycemic profile including controlled blood glucose and improved insulin secretion as previously reported [32]. Therefore, in our work, the antioxidant potential of NRN was investigated not only for improving antioxidant enzyme capacity but also for regulating any increase in the level of ROS. For this purpose, the serum level of NO was assessed before and after treatment. In addition, in chronic diseases, the progression may be partially due to generation of RNS [16]. Altered serum NO levels in DM have been detected by many investigators in prior works with controversial fallouts and consequences [17,18]. Interestingly, we found that hyperglycemia was accompanied with decreased level of antioxidant enzyme superoxide dismutase (SOD) and increase in NO level (Fig 5A and B). Previously, augmented levels of serum NO in patients with DM are not only seen liable for inducing or manipulating DM but have also considered part of diabetes related complications. Unbalanced breakdown of NO may have a role in the pathogenesis of DM, and reduction of nitrogen free radicals because of elevated serum NO could be of value for the evading complications in vascular system associated with DM [19]. Similarly, the proinflammatory mediators IL-6 and TNF-α were considered as the foremost contributors for overproduction of NO as the decline in the levels of cytokines (Fig. 6 A and B) was accompanied positively with reduction in the level of serum NO after NRN treatment.

Importantly, as in this study not only the level of NO was seen to be increased because of hyperglycemia, the level of antioxidant enzyme SOD was also declined. This direct relationship has also been previously reported elsewhere (Meza et al 2019) where activation of vascular endothelial for production of NO because of hyperglycemia has been considered a major factor for production of free reactive species.

It has been concluded that hyperglycemia may be responsible for the increased oxidative stress owing to the increase in level of NO and reduction of SOD. NRN did not only ameliorate the hyperglycemia but also improved the insulin secretion. Further studies should also be done to confirm molecular pathway for hyperglycemia-related increased NO level and reduction in antioxidant enzyme for future improvement in prognostic and therapeutic interventions. This work for the first time shows that naringenin constrains inflammatory mediators like TNF-α and IL-6 further regulating NO production and improvement in antioxidant levels of SOD. This all improves the compromised status of glycemia, lipidemia and altered lever and kidney function via reduction in inflammation and NO-mediated oxidative stress. Consequently, signifying that naringenin is a favorable therapeutic approach as antioxidant and anti-inflammatory natural component, demanding additional investigation in metabolic disorders, as well as in other remedial conditions where such pathophysiological variations are apparent.

## Ethics approval and consent to participate

All applied procedures were approved by the Institutional Biosafety Committee (DGS/3601-04) of University of Agriculture, Faisalabad, Pakistan.

## Consent for publication

N/A

## Availability of supporting data

N/A

## Competing interests

The authors declare that they do not have any conflict of interest for this article.

## Funding

The authors declare that they did not receive any financial support for this paper.

## Authors’ contributions

KR and EA designed study and experimental protocols, searched literature and performed analysis and assessed the preparation of manuscript. IIK carried out the experiments, assisted with data analysis. MAB helped with the manuscript preparation and experiments. MSHA assessed the design of the study, performed statistical analysis and wrote the final draft of manuscript. EA designed study and experimental protocols, searched literature. All the listed authors have read and approved the submitted manuscript.

